## Supplementary Materials for "Ly6G^+^Granulocytes-derived IL-17 limits protective host responses and promotes tuberculosis pathogenesis"

Sharma, P. et al.

**Supplementary Tables: Tables S1-S4**

Table S1: Differentially expressed genes (DEGs) in PMNs identified between each pairwise comparison.

Table S2: Differentially regulated genes in Ly6G$^+$Gra from unvaccinated, uninfected versus BCG-vaccinated, uninfected animals.

Table S3: Gene-set co-regulation analysis of the DEGs with GO-BP in PMNs among the Uninfected control group, *Mtb*-infected group and BCG-vaccinated *Mtb*-infected group.

Tables S4: Gene-set co-regulation analysis of the DEGs with KEGG pathways in PMNs among the Uninfected control group, *Mtb*-infected group and BCG-vaccinated *Mtb*-infected group.

### Supplementary Figure Legends: Figure S1-S4

Supplementary Figure S1

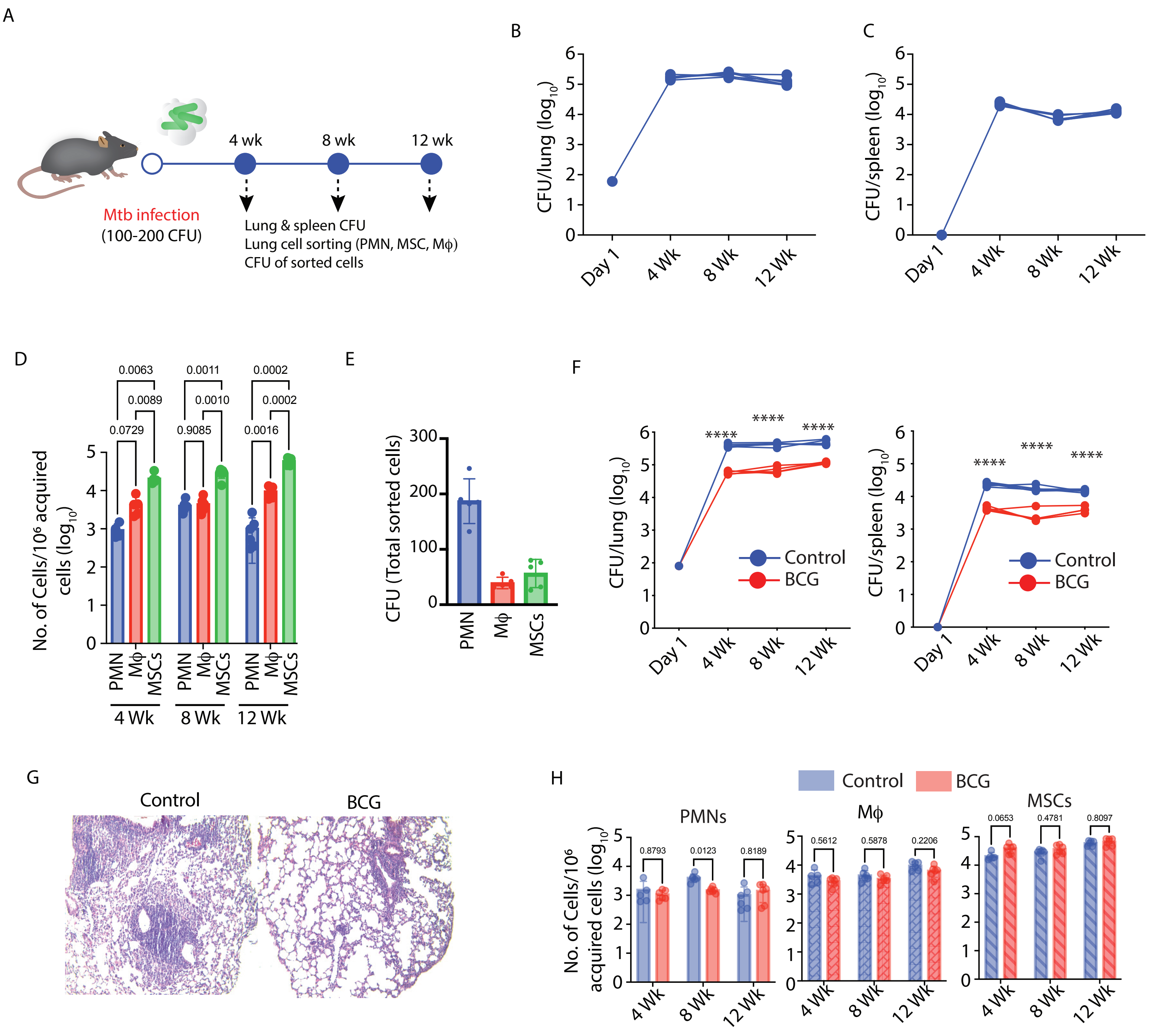

**Figure S1: PMNs harbour a substantial amount of *Mtb*.** (A) Schematic representation of the study design and experimental timeline. C57BL/6 mice were infected with H37Rv, and at 4,8,12 weeks post-infection, mice were sacrificed for various studies. Bacterial burden in (B) Lung and (C) Spleen of infected mice at 4,8,12 weeks post *Mtb* infection. (D) Bar graphs showing the numbers of sorted lung cells (PMNs, Macrophages and MSCs), normalized to per 1 million of total acquired cells at 4,8,12 weeks post-infection. (E) Representative absolute CFU count present inside total sorted PMNs, macrophages and MSCs from the lung. Experimental groups were evaluated using a 2-way ANOVA and 1-way ANOVA statistical test, respectively. Sample size (n)= 5 mice/group in all mice groups. (F) Time course of the lung and spleen bacterial burden in unvaccinated and BCG vaccinated groups (unpaired student's t-test was applied to compare both experimental groups at different time points. (G) Representative H&E-stained lung sections showed less pathology in BCG vaccinated group compared to the unvaccinated group. (H) Quantification of the sorted cells normalized to per 1 million of total acquired cells from the lungs of vaccinated and unvaccinated mice. Experimental groups were evaluated using a 2-way ANOVA statistical test, n = 5 mice/group in both groups.

Supplementary Figure S2

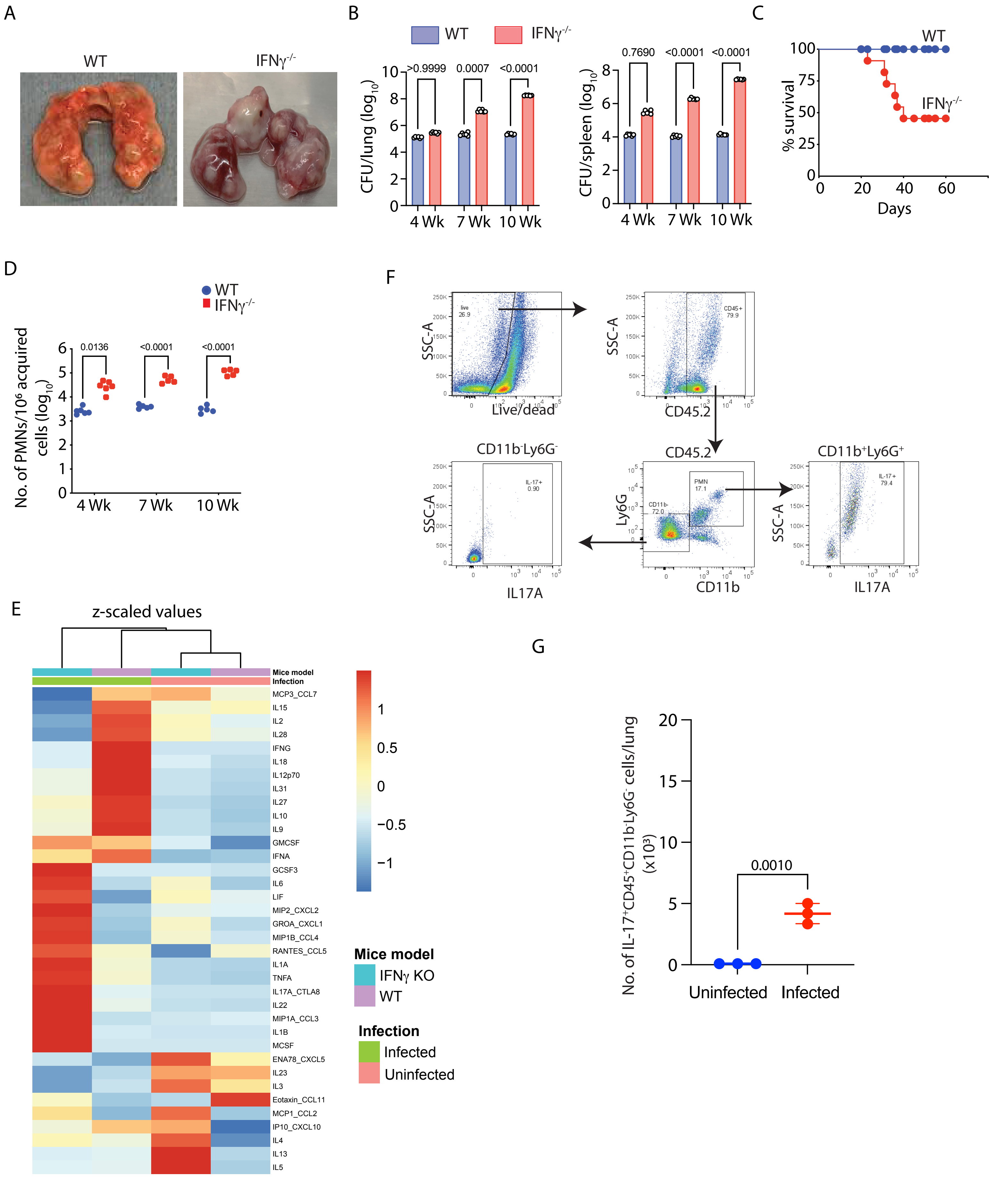

**Figure S2: Contribution of neutrophilia to the severity of TB disease.**

WT and *IFNγ*$^{-/-}$mice were infected with H37Rv, 100 CFU, via the aerosol route. Mice from both groups were euthanized at 4,7- and 10 weeks post-infection time-point. Lung and spleen tissues were harvested and processed for diverse assays, as shown in the study design in Figure 2A. (A) Lung pictures from *Mtb*-infected WT and *IFNγ*$^{-/-}$ mice showing the disease pathology. (B) Enumeration of lung and spleen bacterial *burden* in WT and *IFNγ*$^{-/-}$ mice, 4,7- and 10-weeks post-infection, n= 5-6 mice per experimental group. A 2-way ANOVA statistical test was applied to compare lung and spleen CFU all in WT and *IFNγ*$^{-/-}$ at different time points post-infection. (C) Survival curve of WT and *IFNγ*$^{-/-}$ following *Mtb* infection. Both the strains of mice were infected with 100-200 CFU of H37Rv through an aerosol route followed by incubation till their natural death. The time of death of each mouse was noted and plotted for the survival analysis (n = 12 mice/group). (D) Count of PMNs, normalized to 1 million of total acquired cells, in the lungs of infected WT and *IFNγ*$^{-/-}$ mice at 4-, 7- and 10 weeks post-infection. n= 5-6 mice per experimental group. 2-way ANOVA statistical test was applied to compare groups at different time points post-infection (E) Luminex data: A panel of 36 cytokines and chemokines were quantified from the lung supernatant of uninfected and *Mtb*-infected WT and *IFNγ*$^{-/-}$ mice. All the upregulated and downregulated cytokines and chemokines were compared pairwise between two treatment groups and plotted in the form of a heat map. (F) Lung cells from uninfected WT mice were stained with different antibodies and assayed through flow cytometry to elucidate the cellular source and levels of IL-17 in uninfected mice. Plots showing pooled samples from three mice. (G) Number of CD11b$^{-}$ Ly6G$^{-}$ IL-17$^{+}$ granulocytes in the lungs of uninfected and *Mtb*-infected mice.

Supplementary Figure S3

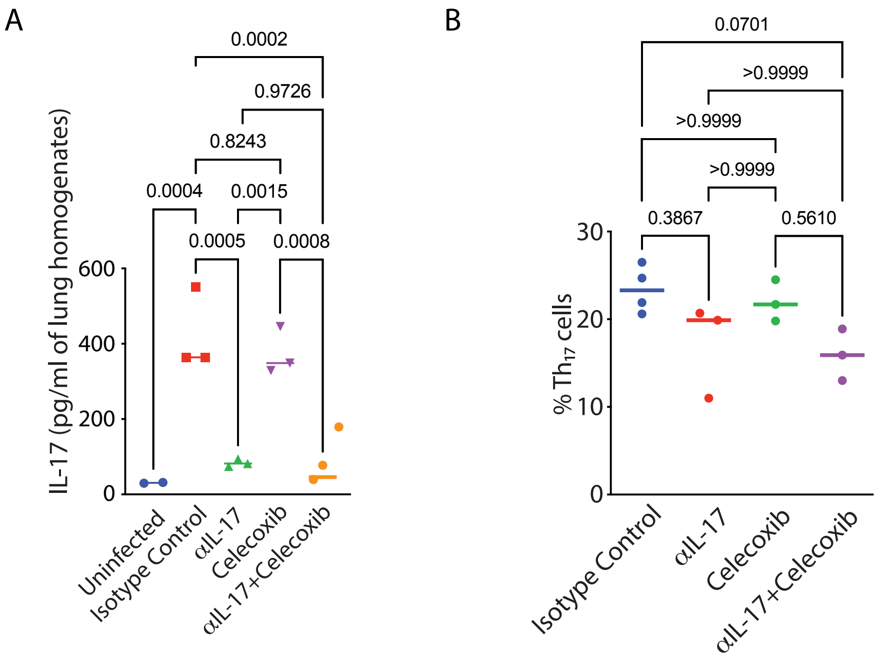

**Figure S3: IL-17 neutralization synergize with COX-2 inhibition and controls TB disease in *IFNγ$^{-/-}$* mice.**

Study plan (as shown in figure 4A): *IFNγ$^{-/-}$* C57BL/6 mice infected with H37Rv at a dose of 100 CFU via the aerosol route. One arm of the *Mtb*-infected mice received an αIL-17 neutralizing antibody (100µg/mouse), and the other arm received an isotype control antibody (100µg/mouse) intraperitoneally twice in a week, starting a day before *Mtb* infection. After 3 weeks of infection, both the groups of mice were further divided into two groups, one that received celecoxib (50mg/kg, orally) for 3 weeks and the other that received vehicle control (DMSO here). Mice from all four groups of mice were euthanized at 6 weeks post *Mtb* infection, and their lungs and spleen were harvested, and serum was stored for ELISA. (A) Lung IL-17 levels in all four mice treatment groups as explained above along with uninfected control. (B) %$Th_{17}$ cells (gated on $CD4^+$ T cells) in the lungs of mice from the same treatment groups described above. The statistical significance of the treatment groups was assessed by applying a one-way ANOVA test. n = 3-4 mice/group.

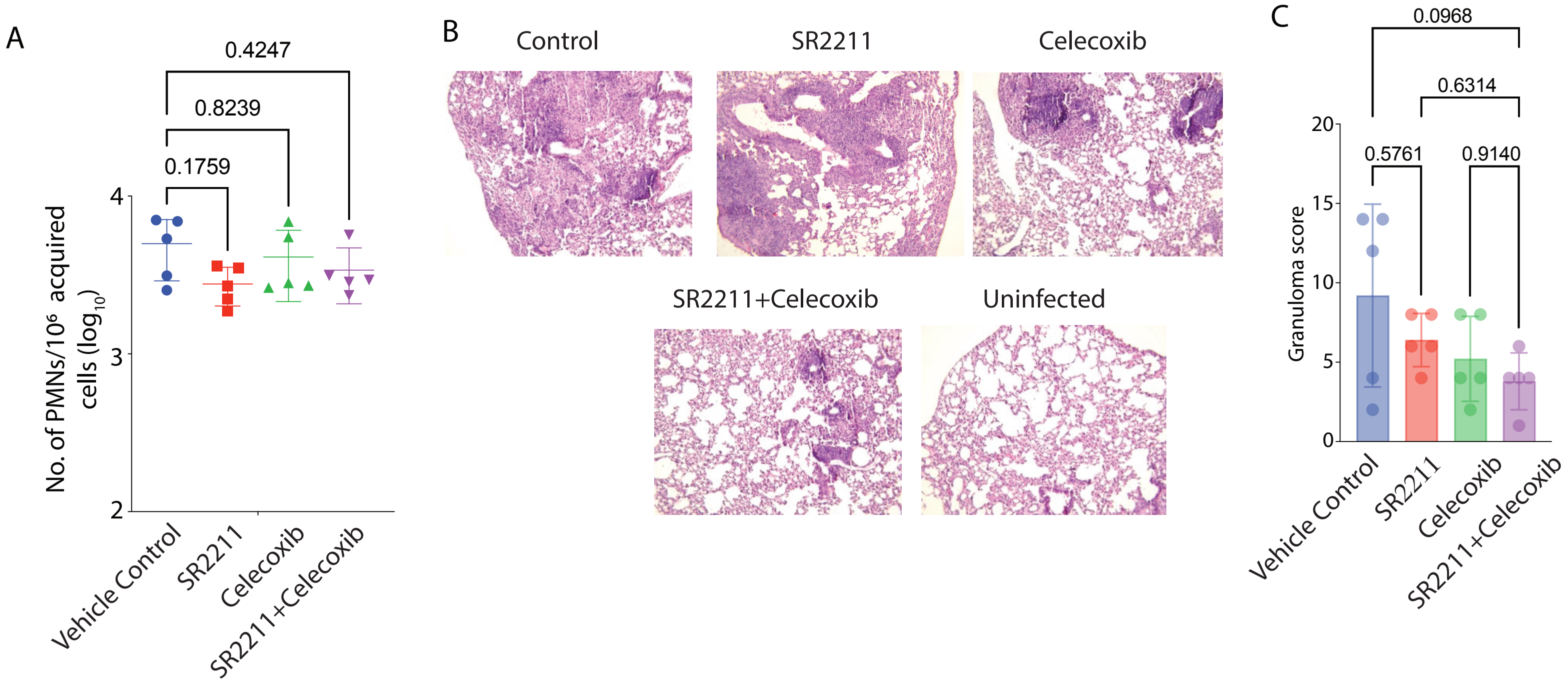
A
0.4247
0.8239
0.1759
No. of PMNs/10⁶ acquired cells (log10)
4
3
2
Vehicle Control
SR2211
Celecoxib
SR2211+Celecoxib
B
Control
SR2211
Celecoxib
SR2211+Celecoxib
Uninfected
C
0.0968
0.6314
0.5761
0.9140
Granuloma score
20
15
10
5
0
Vehicle Control
SR2211
Celecoxib
SR2211+Celecoxib

**Figure S4: IL-17 production inhibition is key to limiting TB severity in WT mice.**

Experimental plan is shown is figure 5(B). C57BL/6 mice were infected with H37Rv at a 100-200 CFU dose via the aerosol route. At 3 weeks post-infection, treatment with celecoxib and SR 2211 (inverse agonist for RORγt) was started for further 3 weeks. Celecoxib was administered orally once a day, and SR 2211 was administered intraperitoneally thrice in a week. At 6 weeks post-infection, mice from the treatment and untreated control groups were sacrificed, and lung and spleen were harvested for CFU, histopathology and FACS analysis. (A) Lung PMN's count normalized to 1 million of total acquired cells in the above-stated treatment groups of mice. The statistical significance of the treatment groups was assessed by applying a one-way ANOVA test, n=5 mice/group. (B) H&E-stained lung sections showing pathology in SR2211+Celecoxib treatment group. (C) Granuloma scoring depicting lung pathology in 4 treatment groups of mice. n= 5 mice per treatment group.
